# Neural, functional, and aesthetic impacts of spatially heterogeneous flicker: A potential role of natural flicker

**DOI:** 10.1101/675090

**Authors:** Melisa Menceloglu, Marcia Grabowecky, Satoru Suzuki

## Abstract

Spatially heterogeneous flicker, characterized by probabilistic and locally independent luminance modulations, abounds in nature. It is generated by flames, water surfaces, rustling leaves, and so on, and it is pleasant to the senses. It affords spatiotemporal multistability that allows sensory activation conforming to the biases of the visual system, thereby generating the perception of spontaneous motion and likely facilitating the calibration of motion detectors. One may thus hypothesize that spatially heterogeneous flicker might potentially provide restoring stimuli to the visual system that engage fluent (requiring minimal top-down control) and self-calibrating processes. Here, we present some converging behavioral and electrophysiological evidence consistent with this idea. Spatially heterogeneous (multistable) flicker (relative to controls matched in temporal statistics) reduced posterior EEG (electroencephalography) beta power implicated in long-range neural interactions that impose top-down influences on sensory processing. Further, the degree of spatiotemporal multistability, the amount of posterior beta-power reduction, and the aesthetic responses to flicker were closely associated. These results are consistent with the idea that the pleasantness of natural flicker may derive from its spatiotemporal multistability that affords fluent and self-calibrating visual processing.

## Introduction

Flickering flames, rippling water surfaces, rustling leaves, and spattering rain drops are pleasant to the senses. Naturally occurring visual flicker such as these is spatially heterogeneous, characterized by probabilistic and locally independent dynamics of luminance modulations.

A prominent perceptual characteristic of spatially heterogeneous flicker is its spatiotemporal multistability. To the extent that local luminance changes are probabilistic and spatially independent, they contain comparable motion energies in multiple directions, speeds, and spatial scales so that spontaneous motion can be seen according to the biases of motion detectors. For example, if leftward-tuned motion detectors happened to be more sensitive (less adapted) than those tuned to other directions, they would be most strongly activated so that a leftward motion would be seen in that region until the sensitive leftward-tuned motion detectors become adapted to be equivalently sensitive to motion detectors tuned to other directions (supported by the literature on flicker motion aftereffects; e.g., Nishida & Ashida, 2000; Mather et al., 2008). Thus, multistable flicker calibrates motion detectors in the sense that more sensitive detectors are more strongly activated than less sensitive ones, thereby adaptively balancing sensitivities across motion detectors tuned to different spatiotemporal patterns. For these reasons, spatially heterogeneous flicker is special in two ways, (1) allowing the perception of spontaneous motion that conforms to the current biases of motion detectors, and (2) calibrating the sensitivities of motion detectors.

Dynamic visual signals often need to be spatially and/or feature-wise integrated to extract movements of behaviorally relevant surfaces and objects. Such integration processes entail long-range neural interactions involving feedback, reflected in the alpha/beta-band power of electrophysiological activity (e.g., Donner & Siegel, 2011; Bastos et al., 2015; Michalareas et al., 2016). For example, MEG beta power in posterior/central scalp regions was increased when stimuli needed to be scrutinized with attention for the extraction of task-relevant information (Donner et al., 2007; Siegel et al., 2008), or when visual signals were spatially integrated to be perceived as a coherently moving object (Aissani et al., 2014). Conversely, posterior/central beta power is reduced when goal-dependent scrutiny that imposes constraints on visual processing is minimal and visual processes transpire according to the current biases of the visual system. For example, EEG posterior/central alpha/beta power was reduced while viewing motions that conformed to biological constraints (Meirovitch et al., 2015), consistent with evidence that the visual system is predisposed to processing biological motion (e.g., Allison et al., 2000; Grossman et al., 2000; Plass et al., 2014). Posterior EEG beta power was reduced while experiencing binocular rivalry (Piantoni et al., 2010), that is, when competing percepts alternated according to changes in the adaptive state of the visual system (e.g., Kim et al., 2006; Wilson, 2007; Alais et al., 2010). Posterior EEG beta power was reduced when an illusory reversed motion was perceived (Piantoni et al., 2010), that is, when motion perception was overtaken by the adaptive biases built up against the motion detectors responding to the veridical motion direction. Although different explanations for alpha/beta-power reductions have been offered by the authors of these studies (see Discussion), a common thread across these findings is that posterior/central alpha/beta-power reductions may reflect visual processes that transpire according to the current biases of the visual system, thereby engaging minimal top-down controls that involve long-range neural interactions.

If this line of reasoning is correct, spatially heterogeneous flicker, whose spatiotemporal multistability allows the activation of motion detectors according to their sensitivity biases, should also reduce central/posterior alpha/beta power. Further, if spatially heterogeneous flicker in nature is pleasant to the senses because the reward system prioritizes the benefit of multistability-based sensory calibration, the degree of multistability afforded by flicker, the amount of central/posterior alpha/beta-power reductions, and the aesthetic preferences for the flicker, should be closely associated. We tested these predictions using flicker displays that presented different degrees of spatiotemporal multistability.

## Experiment 1

Observers viewed a 4-by-4 array of rectangles (e.g., Figure 1) that alternated luminance between high and low values. The sequences of luminance changes were stochastic with specific average rates, generated with a stationary Poisson process where the temporal probability of luminance change (per monitor refresh cycle), ***p_change_***, remained constant. The average flicker rate, 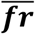, was proportional to ***p_change_***,

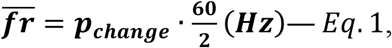

**Figure 1.**
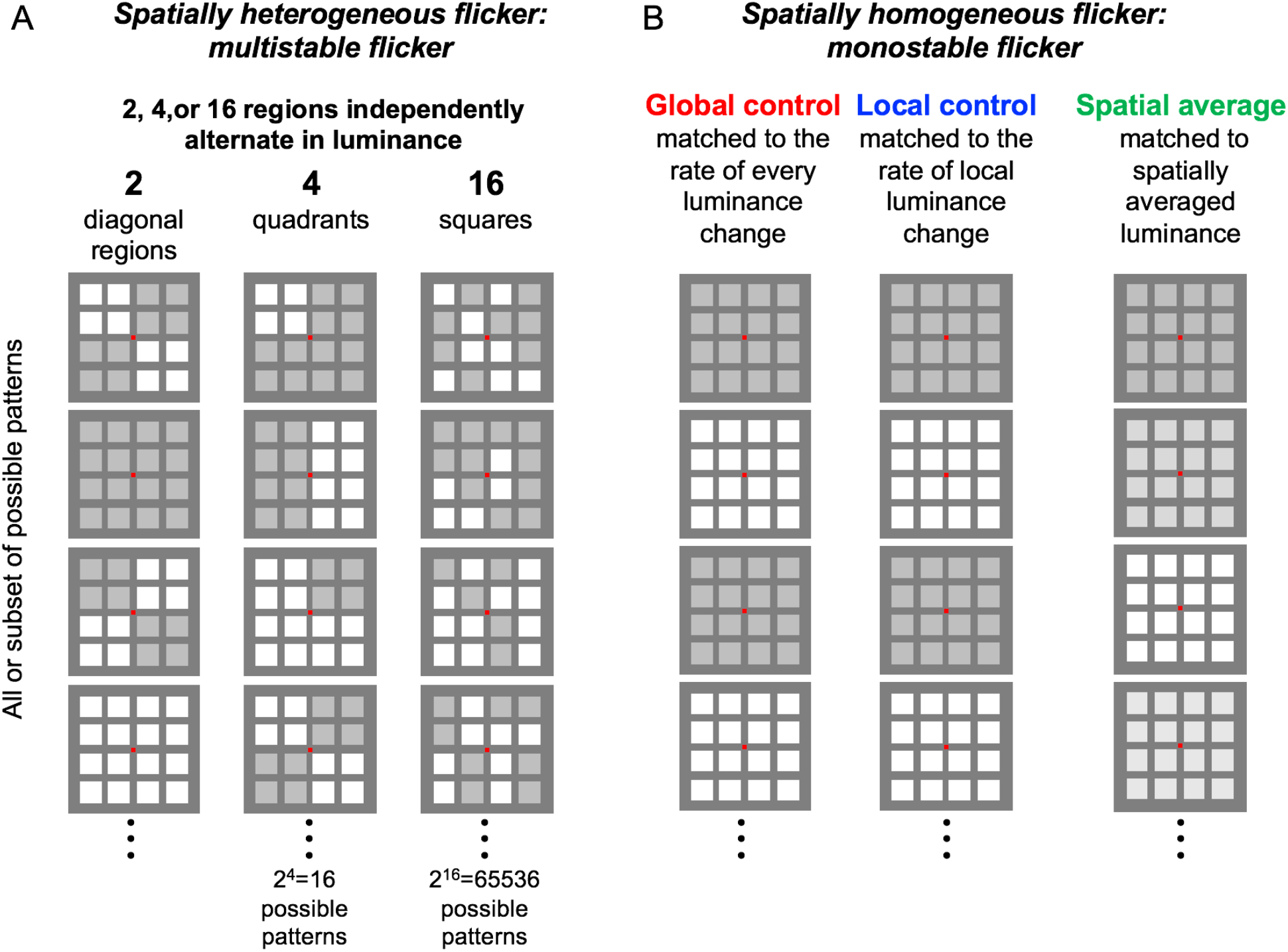
A schematic illustration of the flickering visual stimuli. One, two, four, or sixteen regions independently changed luminance between light and dark values via a Poisson process, with a given probability of luminance change per display refresh cycle (60Hz), generating probabilistic flicker with the temporal Fourier-amplitude profile resembling 1/*f*. **A.** Spatially heterogeneous—*multistable*—flicker (with 2, 4, or 16 regions independently alternating in luminance) that generated the perception of spontaneous motion, with the 16-region condition generating particularly complex motion patterns. B. Spatially homogeneous— *monostabìe*—flicker that did not generate any perception of spontaneous motion, that matched the spatially heterogeneous flicker in terms of (1) the global probability of luminance change (i.e., the probability with which any region may change luminance per display refresh cycle)—the *global control* (all experiments), (2) the local probability of luminance change—the *local control* (all experiments), or (3) the spatially averaged luminance—the *spatial average* (Experiment 2 only). The global- and local-control conditions were included to assess the spatiotemporal effects of multistable flicker over and above the effects of temporal statistics, whereas the spatial-average condition was included to assess the potential effects of low-spatial-frequency representations (see Experiment 2). Each flicker was presented for 5 seconds and observers indicated their aesthetic preference (1-”strongly dislike” to 4-”strongly like”).

where 60 is the display monitor refresh rate and the division by 2 accounts for the fact that a pair of dark and light periods constitutes a luminance-modulation cycle. This method generated flicker with spectral amplitude profiles approximating 1/*f* (except that the maximum flicker rate was 30Hz due to the use of a 60Hz display monitor). We chose this flicker-generating method partly because foveal luminance changes during a typical human experience follow a 1/*f* profile (van Hateren & van der Schaaf, 1996) and partly because 1/*f* flicker tends to be perceived as “less uncomfortable” than flicker with profiles deviating from 1/*f* (Yoshimoto et al., 2017).

We introduced different degrees of spatial heterogeneity by varying the number of independently flickering regions among 1, 2, 4, and 16. In the 1-region condition, all sixteen rectangles synchronously changed their luminance, generating no spatial heterogeneity (Figure 1B). This controlled for the effects of temporal dynamics (see below for details). Spatial heterogeneity was introduced in the 2-, 4-, and 16-region conditions (Figure 1A). In the 2-region condition, the eight rectangles within each pair of diagonal quadrants changed their luminance synchronously, while each pair changed its luminance independently with two parallel Poisson processes with the same ***p_change_***. In the 4-region condition, the four rectangles within each quadrant changed their luminance synchronously, while each quadrant changed its luminance independently, with four parallel Poisson processes with the same ***p_change_***. In the 16-region condition, all sixteen rectangles changed their luminance independently, with sixteen parallel Poisson processes with the same ***p_change_***. The spatiotemporal variety of motion energies in direction, speed, and scale was substantially greater in the 16-region condition relative to the 2- and 4-region conditions, with no spatiotemporal variety in the 1-region control (which can only appear as uniform flicker). Accordingly, the degree of spatiotemporal multistability of perceived motion was substantially greater in the 16-region condition relative to the 2- and 4-region conditions, and was minimal in the 1-region control.

Two aspects of this stimulus design are noteworthy. First, in characterizing the perceptual and neural processing of motion, the complexity of spatiotemporal flicker dynamics can be defined as the number of possible motion interpretations afforded by the flicker. In this sense, the spatiotemporal multistability and complexity of flicker are interchangeable. Second, because the 16 rectangles were always the spatial units of luminance modulation, the dominant spatial frequency composition was similar across all flicker conditions.

To examine the effects of spatiotemporal multistability over and above the effects of its inherent temporal dynamics, we included two types of 1-region flicker stimuli; these were monostable in the sense that they afforded only one spatiotemporal interpretation (uniform flicker). These control stimuli presented two characteristic temporal dynamics inherent in the multistable flicker stimuli (Figure 1B). One was the local luminance-change dynamics of each rectangle, which was determined by ***p_change_*** (Eq.1). Thus, we included the *local-control* stimuli that homogeneously flickered via a Poisson process with ***p_change_***. The other characteristic temporal dynamics was the “global” dynamics reflecting luminance changes across all 16 rectangles, which was determined by the probability with which any rectangle changed luminance (per monitor refresh cycle), ***p_global_***, that increased with the number of independently flickering regions as,

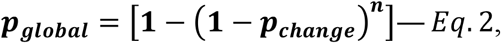

where *n* is the number of independently flickering regions. Thus, we included the *global-control* stimuli that homogeneously flickered via a Poisson process with ***p_global_***. Because we define the degree of flicker multistability as the number of independently flickering regions, these control stimuli had the degree of multistability equaling 1.

Observers viewed the multistable, local-control, and global-control flicker stimuli and rated each one for its aesthetic quality, while their EEG activity was recorded using 64 scalp electrodes. By comparing posterior EEG beta power and aesthetic ratings in response to the multistable flicker stimuli with those in response to the local-control and global-control flicker stimuli, we aimed to identify the effects of viewing spatially heterogeneous—multistable— flicker over and above the effects of viewing spatially homogeneous—monostable—flicker with equivalent temporal statistics. In particular, we considered the broad hypothesis that viewing spatially heterogeneous flicker in nature may be pleasing because its spatiotemporal multistability allows the fluent activation of motion detectors according to their sensitivity biases, which may be prioritized by the reward system because of the potential benefit of the ensuing calibration of the visual motion system. With regard to the current study, this general hypothesis predicted that viewing flicker with a greater degree of multistability should induce lower posterior beta power and higher aesthetic responses (see the introduction and general discussion sections).

### Methods

#### Observers

Twenty-four Northwestern University students gave informed consent to participate for monetary compensation ($10/hr). All were right-handed, had normal hearing and normal or corrected-to-normal vision, and had no history of concussion. They were tested individually in a dimly lit room. Data from six observers were excluded from the analysis due to excessive EEG artifacts (see below). The final sample included 18 observers (7 females) between ages 18 and 29 years (*M* = 22.17, *SD* = 3.33).

#### Stimuli

Each visual display consisted of a small red central-fixation rectangle (0.52° by 0.43°, 21 cd/m^2^) surrounded by a 4-by-4 array of 16 rectangles (12.98° by 10.73° for the overall array), presented against a dark gray background (59 cd/m^2^) (Figure 1). The rectangles (2.60° by 2.15°) were separated horizontally by 3.46° and vertically by 2.86° (center-to-center) from one another. The luminance of each rectangle alternated between the maximum (80 cd/m^2^) and minimum (65 cd/m^2^) values. Visual stimuli were displayed on a 20-in LCD color monitor with 1600-by-1200 pixel resolution at a refresh rate of 60Hz using MATLAB (Version R2016b) with the Psychtoolbox extension (Kleiner et al., 2007) running on a computer with Windows 7. The viewing distance was 90 cm.

The value of ***p_change_*** (the probability of local luminance change per display refresh cycle) was varied between 0.067=4/60 (4 changes per second on average), 0.15=9/60 (9 changes per second on average), and 0.3=18/60 (18 changes per second on average), so that the average flicker rate varied among 2Hz, 4.5Hz, and 9Hz. We included this temporal variation to facilitate the generalizability of the current results; however, we did not have a sufficient number of trials per ***p_change_*** to include it as a factor in the analysis.

On half of the trials, each flicker stimulus was shown with no accompanying sounds—the *visual* trials. On the remaining trials, each flicker stimulus was accompanied by a click sound (a 1 ms white noise burst, 60 dB-SPA, presented through a loud speaker placed behind the display monitor) synchronized with each luminance change—the *audiovisual* trials. This manipulation was included for several reasons. Because spatially heterogeneous flicker in nature is pleasant to look at, any effects of multistable flicker on EEG may potentially be mediated by relaxation or reduced arousal. If so, we would expect any multistable-related EEG effects to disappear or be attenuated on the audiovisual trials with arousing click sounds. In contrast, if multistability-related EEG effects reflect visual processing, they should be equivalent on the visual and audiovisual trials.

We kept the number of trials relatively low partly because we sought robust effects of low intrinsic variability and partly because averaging the beta power across 4 seconds on each trial helped to reduce inter-trial variability. During each block of trials, the 2-region, 4-region, and 16-region multistable flicker (Figure 1A) were shown three times each (once with each of the three values of ***p_change_***), along with the same number of global-control stimuli matched to each multistable flicker in terms of the probability of luminance change at any location (the left panel in Figure 1B), and the local-control stimuli (shown twice with each of the three values of ***p_change_***; the middle panel in Figure 1B), totaling 24 trials. These 24 trials were repeated with the click sounds. Thus, each block consisted of 48 trials, with the trial order randomized per block per observer. Each observer ran two blocks for a total of 96 trials.

#### Behavioral procedures

Each trial began with the appearance of the central fixation marker against a dark gray background. Observers were instructed to fixate the central marker throughout each trial while refraining from blinking. After 1 second of the fixation screen, the flickering squares were displayed for 5 seconds followed by the fixation screen. Observers rated the aesthetic quality of the flicker on the scale from 1-strongly dislike, 2-dislike, 3-like, to 4-strongly like, using a key pad with the correspondingly labelled buttons. Upon key press, the screen turned blank for 1 second during which observers were encouraged to blink. The fixation marker then returned to signal the beginning of the next trial.

#### EEG recording procedures, artifact rejection, and preprocessing

The EEG data were recorded with a sampling rate of 512Hz from 64 scalp electrodes, using a BioSemi ActiveTwo system (see www.biosemi.com for details). Electrooculographic (EOG) activity was monitored using four facial electrodes, one placed lateral to each eye and one placed beneath each eye. Two additional electrodes were placed on the left and right mastoid area. The EEG data were preprocessed using the EEGLAB and ERPLAB toolboxes for MATLAB (Delorme & Makeig, 2004; Lopez-Calderon & Luck, 2014). Data were re-referenced offline to the average of the two mastoid electrodes, bandpass-filtered at 0.01Hz-80Hz (as our spectral-power analyses focused on frequencies between 2Hz and 55Hz), notch-filtered at 60Hz (to remove power-line noise that affected the EEG data from some of the observers), and rid of blink artifacts using the independent component analysis. The continuous EEG data were then segmented into 5.6-second epochs, with each epoch time-locked to the onset of the flickering display, including a prestimulus period of 0.5 seconds (for baselining). Epochs with artifacts were visually identified and manually removed (11% overall for Experiment 1 and 15% for Experiment 2). To reduce the effects of volume conduction and reference electrode choices, as well as to facilitate data-driven EEG source discrimination, we applied a surface-Laplacian transform to all EEG data (Hjorth, 1980; Kayser and Tenke, 2006; Tenke and Kayser, 2012), using the Perrin et al.’s method (e.g., Perrin et al., 1987; Perrin et al., 1989a; 1989b) with a typical set of parameter values (Cohen, 2014).

#### EEG data analysis

For each trial for each observer, the surface-Laplacian transformed EEG waveform from each scalp site was decomposed into a time series of spectral power using 60 Gabor wavelets with center frequencies *f*’s and the factor *n*’s (proportional to the temporal standard deviation of the wavelet, *n* = 2*πf* · *SD*) that were logarithmically spaced (because neural temporal-frequency tunings tend to be approximately logarithmically scaled; e.g., Hess & Snowden, 1992; Lui et al., 2007). The wavelet center frequencies spanned the range of 2Hz to 55Hz and the *n* values spanned the range of 3 to 16, resulting in temporal resolutions of *SD*=239ms (at 2Hz) to *SD*=46ms (at 55Hz) and spectral resolutions of *FWHM* (full width at half maximum) =1.56Hz (at 2Hz) to *FWHM*=8.09Hz (at 55Hz). These values struck a good balance for the temporal-spectral-resolution trade-off, and are typically used in the literature (e.g., Cohen, 2014).

The time-frequency matrices from the individual trials were averaged per condition per observer. The portion of the time-frequency matrices corresponding to the pre-stimulus baseline period (−0.4 second to −0.15 second relative to flicker onset) were averaged across time, averaged across trials within each condition, and then averaged across all conditions per frequency per observer. The average time-frequency matrix for each condition was divided by this common baseline (per frequency per observer) and was then converted to dB (by taking the base-10 log and multiplying by 10). These values were then averaged across the posterior scalp sites (see the topographic plots in Figure 2B) which were responsive to multistable flicker. All statistical analyses were conducted with observer as the random effect.

**Figure 2.**
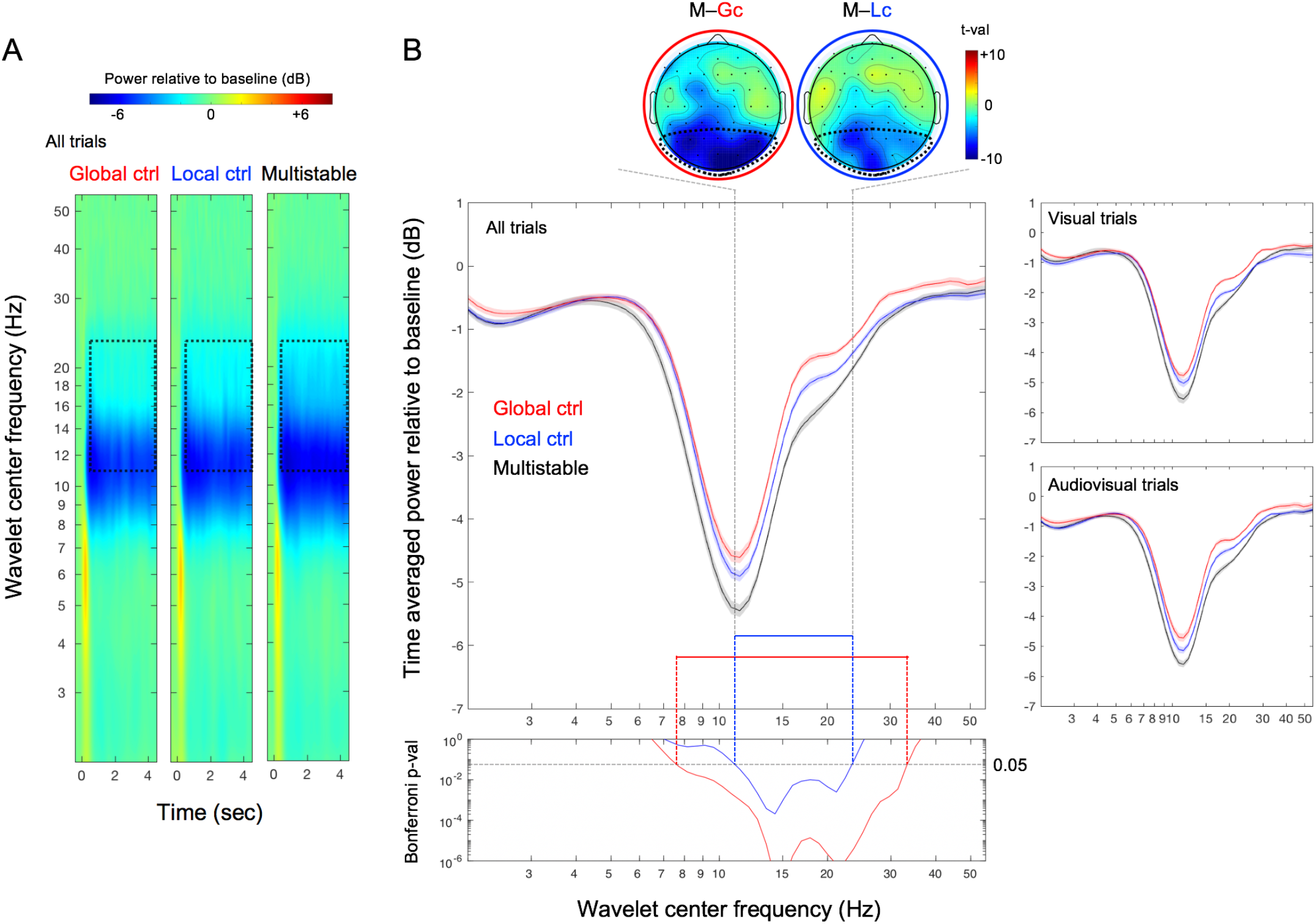
Results of Experiment 1. **A.** EEG spectral power in dB baselined to −0.4 to −0.15 second prestimulus period (shown in color scale, averaged across the posterior scalp sites indicated with the dotted boundaries in the topographic plots shown in B) for wavelet center frequencies from 2Hz to 55Hz as a function of time (relative to flicker onset at 0) for the global-control (left), local-control (middle), and multistable (right) conditions. The dotted rectangles indicate the 0.5 to 4.5 second time period over which EEG spectral power was averaged for analysis and the ~11Hz to ~23Hz frequency range in which EEG spectral power reductions were significantly larger for the multistable condition than for each of the control conditions (see B). **B.** Main panel. Time averaged posterior EEG power (in dB) as a function of wavelet center frequency for the global-control (red), local-control (blue), and multistable (black) conditions; the right panels show the data from the visual and audiovisual trials separately. Bottom panel. Bonferroni-corrected *p*-values for the comparison between the multistable condition and each of the monostable control conditions as a function of wavelet center frequency. The ~11Hz to ~23Hz frequency range in which EEG power was significantly reduced in the multistable condition relative to both control conditions is indicated by the tall dashed lines, with the two topographic plots at the top showing the posterior focus of this beta-range power reduction (in *t*-values) associated with perceiving multistable flicker. The shaded regions represent ±1 *SEM* adjusted for the repeated-measures comparisons (Morey, 2008).

### Results and discussion

#### Multistable flicker reduces posterior beta power

All flicker stimuli (global control, local control, and multistable) reduced EEG spectral power between ~8Hz and ~25Hz (Figure 2A) in the posterior scalp region where beta-power reductions associated with visual multistability were prominent (topographic plots at the top of Figure 2B). Notably, power reductions in the beta range (~11Hz to ~23Hz) were pronounced in the multistable condition relative to the control conditions (see the regions marked by the dotted rectangles in Figure 2A). This effect is more clearly seen when the EEG power is averaged over the flicker period (0.5–4.5 second; the main panel in Figure 2B). The Bonferroni-corrected *p*-values comparing the multistable condition with each of the control conditions (via pairwise *t*-tests with *df*=17; the bottom panel in Figure 2B) across the 60 wavelet center frequencies show that power reductions within the range of ~11Hz to ~23Hz was significantly larger in the multistable condition than in either of the two control conditions. The topographic plots (at the top of Figure 2B) confirm that these effects were localized within the posterior scalp regions. The multistable condition reduced beta power similarly whether visual flicker was presented alone or accompanied by click sounds (the right panels in Figure 2B). The posterior scalp focus and crossmodal invariance are consistent with the interpretation that the beta-power reductions were visually driven.

#### A greater degree of multistability induces a larger beta-power reduction

How did the posterior beta-power reduction depend on the degree of multistability operationalized as the number of independently flickering regions? We note that the local-control condition can be considered a 1-region “multistable” condition because ***p_change_*** in the local-control condition was matched to ***p_change_*** for each independently flickering region in the multistable conditions. The local-control condition would also be the global-control condition for 1-region flicker because ***p_change_*** per region would be the same as the ***p_global_*** (the probability of luminance change at any region) when the entire array flickered homogeneously. Thus, for analyzing the effects of the degree of multistability, we designate the local-control condition to represent both the multistable and global-control conditions for 1-region flicker.

Three observations stand out (Figure 3A); (1) even the lowest 2-region multistability substantially reduced posterior beta power relative to the monostable (1-region) flicker with matched local temporal statistics (the blue asterisks in Figure 3A), (2) all degrees of multistability reduced beta power relative to the corresponding global-control conditions with matched global temporal statistics (the red asterisks in Figure 3A), and (3) the highest degree of multistability with sixteen independently flickering regions caused the largest reduction in beta power (the black asterisks in Figure 3A). Consistent with the data shown in Figure 2A, the multistable conditions similarly reduced beta power with or without the accompanying click sounds (the right panels in Figure 3A), suggesting that the beta-power reductions associated with viewing multistable displays reflect visual processing rather than changes in arousal.

**Figure 3.**
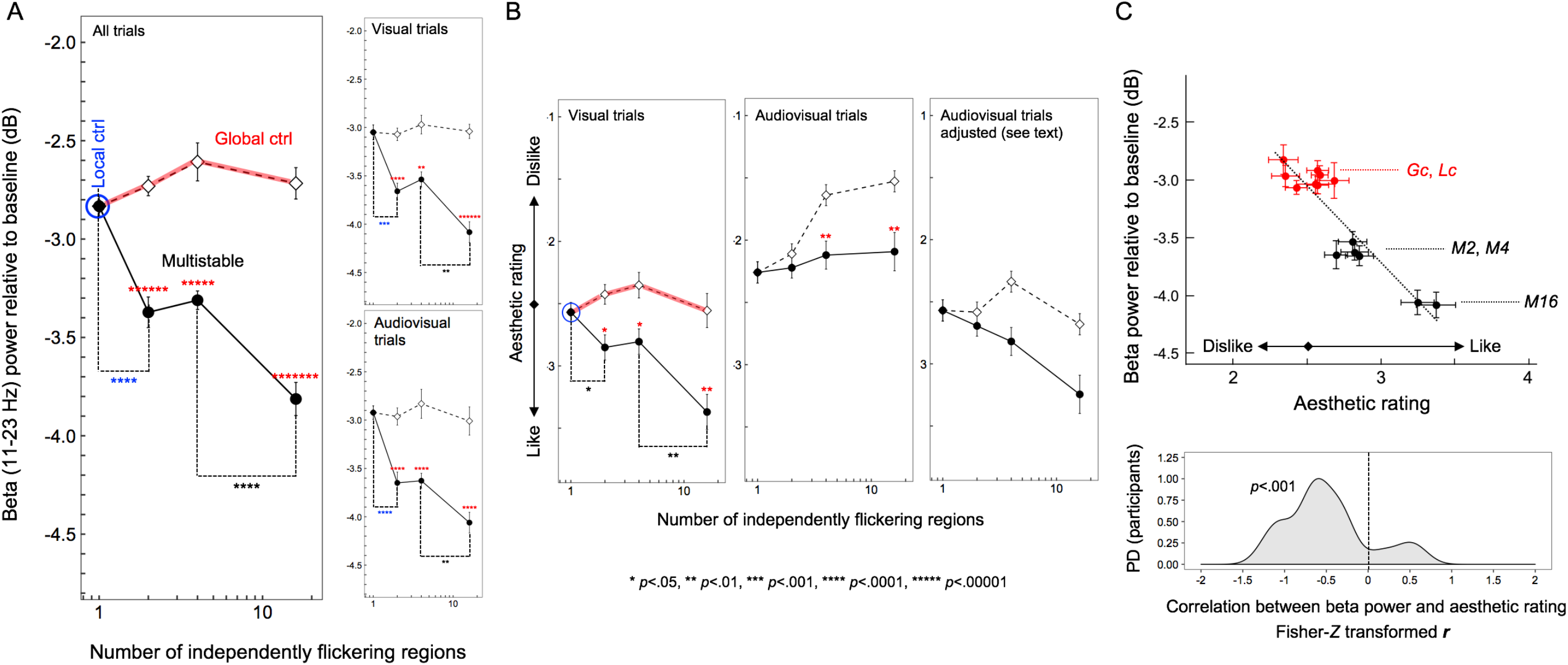
Results of Experiment 1. **A.** Time-averaged posterior EEG beta power (~11Hz to ~23Hz for which EEG power was significantly reduced for the multistable condition relative to all control conditions; see Figure 2) shown as a function of the number of independently flickering regions. Note that the local-control condition can be considered a “multistable” condition with the number of independently flickering regions being 1 because the rate of flicker was matched to the rate at which each region was independently flickered in the multistable condition. Overall, the multistable condition for all degrees of multistability (with 2, 4, or 16 independently flickering regions) significantly reduced beta power relative to the corresponding global-control conditions (red asterisks); the lowest degree of multistability (with only 2 independently flickering regions) significantly reduced beta power relative to the local-control condition (blue asterisks); the most complex multistability (with 16 independently flickering regions) significantly reduced beta power relative to the 4-region multistability (black asterisks). The pattern of results was similar when the data were examined separately for the visual and audiovisual trials (right panels). **B.** Similar analyses on aesthetic ratings. When the y-axis is ordered from “strongly like” (4) to “strongly dislike” (1), the pattern of aesthetic ratings for the visual trials (left panel) strongly resembles the pattern of EEG beta-power reductions—the beta-power-reducing multistable conditions being generally preferred and the strongly beta-power-reducing 16-region multistability especially preferred—suggesting that greater beta-power reductions are associated with higher aesthetic ratings. However, unlike the EEG beta-power reductions that were similar between the visual and audiovisual trials (see A), the aesthetic ratings were lowered in the audiovisual trials (middle panel). However, if we make the simple assumption that the synchronized click sounds (which sounded unpleasant to most participants) subtractively lowered aesthetic preferences without interacting with flicker effects, the appropriately adjusted ratings (right panel) resembles those from the visual trials (left panel; see main text for details). **C(upper).** Scatter plot showing the relationship between the posterior EEG beta-power reductions and aesthetic ratings (adjusted to exclude the general sound effects on ratings; see text for details) across the flicker conditions (*Lc*–local control; *Gc*–global control; *M2, M4*, and *M16*–multistable with 2, 4, and 16 independently flickering regions, respectively) with visual and audiovisual presentations. The negative relationship indicates that greater EEG beta-power reductions were associated with higher aesthetic ratings in response to the flicker conditions. **C(lower).** Probability-density distribution (PD) of the Fisher-*Z* transformed *r* (Pearson’s correlation coefficient) values for the above relationship computed for the individual observers; in support of the negative relationship above, the distribution of *r_z_* is substantially shifted in the negative direction. The error bars represent ±1 *SEM* adjusted for repeated-measures comparisons (Morey, 2008).

#### Aesthetic preferences are associated with the amount of beta-power reductions in response to flicker multistability

The aesthetic ratings (coded from “like” to “dislike”) closely mirrored the pattern of beta-power reductions (the left panel in Figure 3B), indicating that higher degrees of multistability generated both larger beta-power reductions and greater aesthetic preferences.

Interestingly, although beta-power reductions were unaffected by the click sounds, aesthetic preferences were lowered by the sounds (the middle panel in Figure 3B). The click sounds were synchronized to luminance changes occurring at any location so that they were more frequent for flicker with a greater degree of multistability (that contained more independently flickering regions). Observers informally indicated that frequent click sounds were unpleasant. Thus, although observers were instructed to rate the visual aspect of flicker while ignoring the sounds, the unpleasant sounds might have pulled the aesthetic ratings down. A simple possibility is that the sounds subtractively lowered aesthetic ratings without interacting with the effects of visual flicker multistability. Consistent with this possibility, if we linearly adjust the ratings from the audiovisual trials so that their condition average (i.e., average across the multistable and global-control conditions) is the same as that for the visual trials for each degree of multistability, that is, if we linearly remove the main effect of click-sound frequency on ratings, the rating pattern from the audiovisual trials become similar to that from the visual trials (the rightmost panel in Figure 3B). This result, combined with the result that the effects of the multistable and control flicker on the posterior beta power were equivalent with or without the click sounds (the right panels in Figure 3A), suggests that the posterior beta-power reduction is selectively associated with the visual component of aesthetic ratings. This interpretation hinges on the similarity between the beta-power-reduction pattern and the aesthetic-rating pattern (subtractively adjusted for the audiovisual trials) as a function of the degree of multistability across the visual trials (the 7 beta-power values in the upper right panel in Figure 3A and the 7 rating values in the leftmost panel in Figure 3B) and the audiovisual trials (the 7 beta-power values in the lower right panel in Figure 3A and the 7 rating values in the rightmost panel in Figure 3B). The similarity is captured in the 14-point scatterplot showing the negative relationship between the posterior beta power and aesthetic rating across the multistable and control conditions with visual and audiovisual presentations (Figure 3C, upper panel). It is clear from the scatter plot that the relationship is driven by the greater beta-power reductions and higher aesthetic ratings for the multistable conditions, especially for the highest degree of multistability (with 16 independently flickering regions). To confirm the reliability of this relationship, we computed the Fisher-*Z* transformed correlation coefficients, *r*_z_’s, for individual observers and plotted the probability density distribution of *r*_z_ (Figure 3C, lower panel). The distribution is substantially shifted in the negative direction, *r*_z_ *mean* = .490, *sem* = .117, *t*_17_ = 4.186, *p*=.00062, suggesting that the posterior beta-power reductions are reliably associated with visual aesthetic responses to flicker multistability.

Taken together, spatially heterogeneous (multistable) flicker reduced posterior EEG beta power (~11Hz to ~23Hz) relative to the spatially homogeneous controls that were matched in the temporal dynamics of local luminance changes (the local control) and luminance changes at any location (the global control). Even the lowest degree of multistability (with only two independently flickering regions) induced a substantial beta-power reduction, but the highest degree of multistability (with sixteen independently flickering regions) induced a much larger beta-power reduction. Beta-power reductions were equivalent with or without the accompanying click sounds and was closely associated with aesthetic responses to flicker multistability. These results suggest that multistable flicker reduces posterior beta power and contributes to aesthetic preferences.

The results presented in Figures 2 and 3 are relatively clean, and are consistent with our general hypothesis. However, the specific pattern of posterior focus and the specific frequency range of the multistability-dependent beta-power reductions, the lack of auditory effects on the posterior beta-power reductions, and the subtractive effect of click sounds on aesthetic ratings, were not anticipated. It was thus necessary to replicate the results of Experiment 1.

## Experiment 2

The design of this experiment was the same as that of Experiment 1 except that an additional condition was included. The visual system internally generates spatially averaged flicker signals through low-spatial-frequency channels (e.g., Stromeyer et al., 1982) while processing spatially heterogeneous flicker. Thus, the posterior beta-power reductions and aesthetic responses we obtained in association with spatiotemporal multistability in Experiment 1 might potentially be driven by the spatially averaged temporal dynamics. Further, even if the posterior beta-power reductions and aesthetic responses were driven by spatiotemporal multistability, given that the low-spatial-frequency channels concurrently generate spatially averaged versions while people experience naturally occurring spatially heterogeneous flicker (e.g., flames, water surfaces) and given that associations can form between neural assemblies on the basis of coincident activations (e.g., Hebb, 1949; Rumelhart & McClelland, 1986), the viewing of a spatially averaged version of multistable flicker may associatively induce posterior beta-power reductions and/or aesthetic responses. In particular, coincidence-based associations have been shown to play a major role in generating visual preferences (e.g., Palmer & Schloss, 2010; Strauss et al., 2013).

We thus included a *spatial-average* condition where all rectangles were assigned the spatially averaged luminance of the corresponding spatially heterogeneous (multistable) flicker at each time point (i.e., each monitor refresh frame). If spatially averaged dynamics rather than spatiotemporal multistability drives the beta-power reductions and aesthetic responses, the effects of the multistable and spatial-average conditions should be equivalent. Regarding the associative activation possibility, any spatial averaging performed by the low-spatial-frequency channels would have limited spatial range, so that some spatiotemporal multistability would still be present in low-spatial-frequency channel outputs. Therefore, any associative induction of posterior beta-power reductions and aesthetic responses by the fully spatially averaged flicker (with no spatial heterogeneity) is expected to be weaker. Thus, we expect that even if spatially averaged dynamics associatively induced posterior beta-power reductions and aesthetic responses, the effects produced by the spatial-average condition should be weaker than those produced by the multistable condition.

### Methods

#### Observers

Twenty-four Northwestern University students gave informed consent to participate for monetary compensation ($10/hr). All observers were right-handed, had normal hearing and normal or corrected-to-normal vision, and had no history of concussion. They were tested individually in a dimly lit room. Data from six observers were excluded from the analysis due to excessive EEG artifacts. The final sample included 18 observers (11 females, 1 nonbinary) between ages 18 and 23 years (*M* = 19.94, *SD* = 1.29). The fact that six observers were excluded from analysis based on EEG artifacts in both experiments was coincidental.

#### Stimuli

The stimuli were the same as those used in Experiment 1 except that trials of the spatial-average condition (presenting spatially averaged versions of the multistable stimuli) were added. Because the spatial-average stimuli matched the multistable stimuli for each degree of multistability (2, 4, or 16 independently flickering regions) as did the global-control stimuli (see Experiment 1), the number of the spatial-average trials was the same as the number of the global-control trials.

#### Behavioral procedures, EEG recording procedures, artifact rejection, and preprocessing, *and* EEG data analysis

These were the same as those in Experiment 1.

### Results and discussion

#### Multistable flicker reduces posterior beta power

As in Experiment 1, the plot of EEG spectral power (color coded) as a function of wavelet center frequency (y axis) and time (x axis) shows that, although all flicker stimuli reduced EEG power between ~8Hz and ~20Hz, power reductions in the beta range (~14Hz to ~20Hz) appear pronounced for the multistable condition relative to the global-control, local-control, and spatial-average conditions (see the regions marked by the dotted rectangles in Figure 4A). This difference is more clearly seen when EEG power is averaged across the flicker period (0.5–4.5 second) (the main panel in Figure 4B). Bonferroni-corrected *p*-values comparing the multistable condition with each of the other conditions (via pairwise *t*-tests with *df*=17) across the 60 wavelet center frequencies (the bottom panel in Figure 4B) show that beta-power reductions were significantly larger for the multistable condition than for any of the other conditions in the range of ~14Hz to ~20Hz, which falls within the beta range identified in Experiment 1. The topographic plots (at the top of Figure 4B) confirm that these effects were localized within the same posterior scalp regions identified in Experiment 1. As in Experiment 1, the multistable condition reduced beta power similarly whether flicker was presented alone or accompanied by click sounds (the right panels in Figure 4B). The posterior scalp focus and crossmodal invariance are again consistent with the interpretation that the beta-power reductions were visually driven.

**Figure 4.**
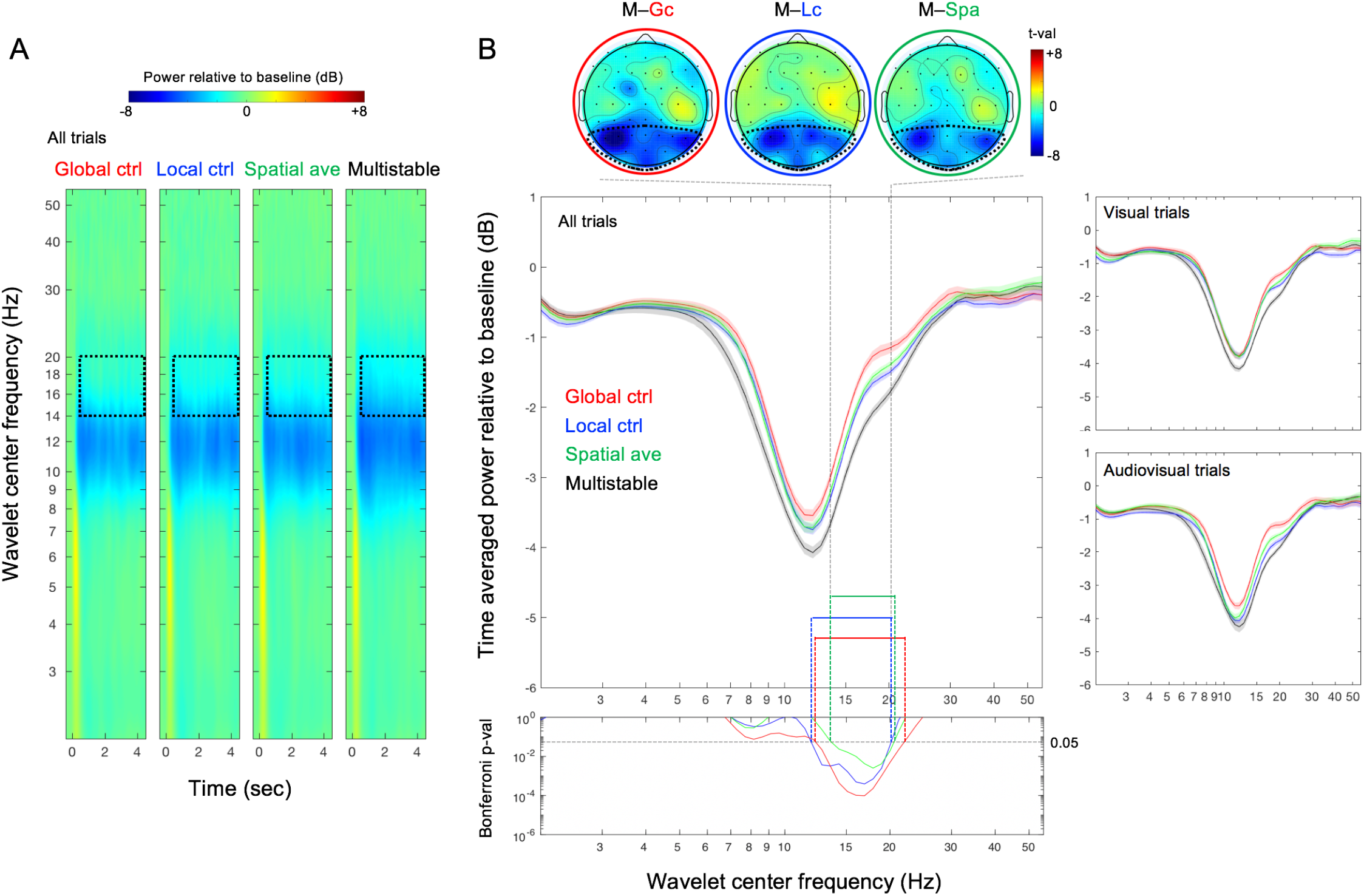
Results of Experiment 2. **A.** EEG spectral power in dB baselined to −0.4 to −0.15 second prestimulus period (shown in color scale, averaged across the posterior channels indicated with the dotted boundaries in the topographic plots shown in B) for wavelet center frequencies from 2Hz to 55Hz as a function of time (relative to flicker onset at 0) for the global-control, local-control, spatial-average, and multistable conditions (from left to right). The dotted rectangles indicate the 0.5 to 4.5 second time period over which EEG spectral power was averaged for analysis and the ~14Hz to ~20Hz frequency range in which EEG spectral power reductions were significantly larger for the multistable condition than for each of the other conditions (see B). **B.** Main panel. Time averaged posterior EEG power (in dB) as a function of wavelet center frequency for the global-control (red), local-control (blue), spatial-average (green), and multistable (black) conditions; the right panels show the data from the visual and audiovisual trials separately. Bottom panel. Bonferroni-corrected *p*-values for the comparison between the multistable condition and each of the other conditions as a function of wavelet center frequency. The ~14Hz to ~20Hz frequency range in which EEG power was significantly reduced in the multistable condition relative to all other conditions is indicated by the tall dashed lines, with the three topographic plots at the top showing the posterior focus of this beta-range power reduction (in *t*-values) associated with perceiving multistable flicker. The shaded regions represent ±1 *SEM* adjusted for the repeated-measures comparisons (Morey, 2008).

#### A greater degree of multistability induces a larger beta-power reduction

As noted in the results section of Experiment 1, for the analysis of the effects of the degree of multistability (operationalized as the number of independently flickering regions), the local-control condition represents the global-control, spatial-average, and multistable conditions for the case of 1-region flicker.

As in Experiment 1, three observations stand out (Figure 5A); (1) even the lowest 2-region multistability substantially reduced beta power relative to the monostable (1-region) flicker with matched local temporal statistics (the blue asterisks in Figure 5A), (2) all degrees of multistability reduced beta power relative to the corresponding global-control conditions with matched global temporal statistics (the red asterisks in Figure 5A), and (3) the highest degree of multistability with sixteen independently flickering regions generated the largest beta-power reduction (the black asterisks in Figure 5A). Furthermore, all degrees of multistability reduced beta power relative to the corresponding spatial-average conditions (the green asterisks in Figure 5A). Consistent with Experiment 1, the multistable conditions reduced beta power similarly with or without the accompanying click sounds (the right panels in Figure 5A), again, suggesting that the beta-power reductions associated with viewing multistable displays reflect visual processing rather than changes in arousal

**Figure 5.**
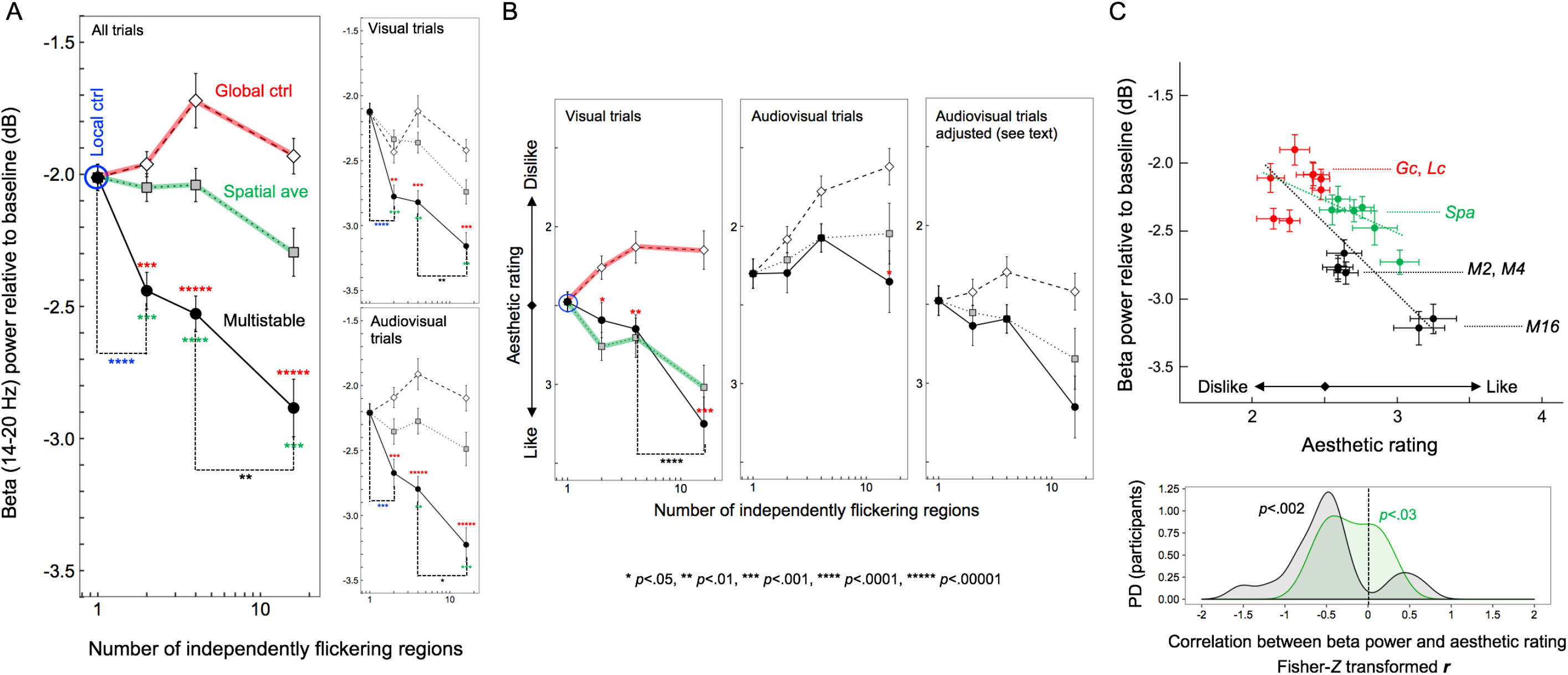
Results of Experiment 2. **A.** Time-averaged posterior EEG beta power (~14Hz to ~20Hz for which EEG power was significantly reduced for the multistable condition relative to all other conditions) shown as a function of the number of independently flickering regions. Note that the local-control condition can be considered a “multistable” condition with the number of independently flickered regions being 1 because the rate of flicker was matched to the rate at which each region was independently flickered in the multistable condition. Overall, the multistable condition for all degrees of multistability (with 2, 4, or 16 independently flickering regions) significantly reduced beta power relative to both the global-control (red asterisks) and spatial-average (green asterisks) conditions; the lowest degree of multistability (with only 2 regions independently flickering) significantly reduced beta power relative to the local-control condition (blue asterisks); the most complex multistability (with 16 regions independently flickering) significantly reduced beta power relative to the 4-region multistability (black asterisks). The pattern of results was similar when the data were examined separately for the visual and audiovisual trials (right panels). **B.** Similar analyses on aesthetic ratings. When the y-axis is ordered from “strongly like” (4) to “strongly dislike” (1), the pattern of aesthetic ratings for the visual trials (left panel) overall resembles the pattern of EEG beta-power reductions—the beta-power-reducing multistable conditions being generally preferred and the strongly beta-power-reducing 16-region multistability especially preferred—suggesting that greater beta-power reductions are associated with higher aesthetic ratings. The multistable and spatial-average conditions were equivalently preferred (left panel), potentially reflecting the associative learning of preference for spatially averaged versions that are internally generated by the low-spatial-frequency channels while multistable flicker is viewed (see text for details). As in Experiment 1 (see Figure 3B), the aesthetic ratings were lowered in the audiovisual trials (middle panel). Consistent with Experiment 1, if we make the simple assumption that the synchronized click sounds (which sounded unpleasant to most observers) subtractively lowered aesthetic preferences without interacting with flicker effects and apply the same linear adjustment we used in Experiment 1, the ratings from the audiovisual trials become similar to those from visual trials (the right panel). **C(upper).** Scatter plot showing the relationship between the posterior EEG beta-power reductions and aesthetic ratings (adjusted to exclude the general sound effect on ratings; see text for details) across the flicker conditions (*Lc*–local control; *Gc*–global control; *Spa*–spatial average; *M2*, *M4*, and *M16*–multistable with 2, 4, and 16 independently flickering regions, respectively) with visual and audiovisual presentations. The negative relationship indicates that greater EEG beta-power reductions were associated with higher aesthetic ratings in response to the flicker conditions; the negative relationship is stronger across the control (red symbols) and multistable (black symbols) conditions (black linear regression fit) than across the control (red symbols) and spatially averaged (green symbols) conditions (green linear regression fit). **C(lower).** Probability-density distributions (PD) of the Fisher-Z transformed *r* (Pearson’s correlation coefficient) values for the above relationships computed for the individual participants; the distribution for the control-multistability relationship is strongly shifted in the negative direction (black area) whereas the shift is moderate for the control-spatial-average relationship (green area). The error bars represent ±1 *SEM* adjusted for repeated-measures comparisons (Morey, 2008).

#### Aesthetic preferences are associated with the amount of beta-power reductions in response to flicker multistability

The aesthetic ratings (coded from “like” to “dislike”) for the multistable and global-control conditions for the visual trails closely mirrored the pattern of beta-power reductions (the red and black curves in the leftmost panel in Figure 5B), indicating that higher degrees of multistability generated both larger beta-power reductions and greater aesthetic preferences, replicating Experiment 1. Also consistent with Experiment 1, although aesthetic ratings were lowered by the click sounds (the middle panel in Figure 5B), linearly removing the main effect of click-sound frequency on ratings (the rightmost panel in Figure 5B) made the rating pattern from the audiovisual trials similar to that from the visual trials (the leftmost panel in Figure 5B). Thus, the converging results from the two experiments suggest that the sounds subtractively lowered the ratings without interacting with the effects of visual flicker.

The unique aspect of this experiment was the inclusion of the spatial-average condition. We considered two possibilities. Beta-power reductions and aesthetic responses might be driven by the spatially averaged temporal dynamics of multistable flicker rather than by the spatiotemporal multistability per se, predicting that the multistable and spatial-average conditions should produce equivalent effects. This possibility was ruled out because the spatial-average conditions were substantially less effective at reducing posterior beta power than the corresponding multistable conditions (Figure 5A), confirming the unique role of spatiotemporal multistability in reducing posterior beta power. The other possibility we considered was that even if spatiotemporal multistability drives the beta-power reduction and aesthetic effects, their spatially averaged versions might still induce a weaker version of those effects through learned associations because spatial-averaged versions (with limited spatial ranges) are always present in the output from the low-spatial-frequency channels. In support of this possibility, though the spatial-average condition did not reliably produce beta-power reductions (Figure 5A), it was equivalently effective to the multistable condition at generating aesthetic responses (Figure 5B). This is consistent with the fact that visual preferences are susceptible to coincidence-based associative learning (e.g., Palmer & Schloss, 2010; Strauss et al., 2013).

Importantly, as in Experiment 1, the pattern of posterior beta-power reductions and the pattern of aesthetic ratings are closely associated as a function of the degree of multistability across the visual and audiovisual trials. This is evidenced by the negative relationship between the 14 beta-power values (the white squares and black circles shown across the upper and lower right panels in Figure 5A) and the 14 aesthetic-rating values (the white squares and black circles shown across the leftmost and rightmost panels in Figure 5B) for the multistable and control conditions. The relationship is displayed as a scatter plot in the upper panel in Figure 5C; see the red and black symbols with the dotted black linear regression line. Consistent with Experiment 1, this relationship is driven by the greater beta-power reductions and higher aesthetic ratings for the multistable conditions, especially for the highest degree of multistability (with 16 independently flickering regions). The scatter plot also shows the relationship for the spatial-average and control conditions; see the red and green symbols with the dotted green linear regression line. The relationship is weaker but still negative, suggesting that the spatially averaged versions of the multistable stimuli moderately induced the neural-behavioral responses that are similar to those induced by the corresponding multistable stimuli.

To evaluate the reliability of these relationships, we computed the Fisher*-Z* transformed correlation coefficients, *r*_z_’s, for individual observers and plotted their probability density distributions (as in Experiment 1). The distribution for the relationship for the multistable and control conditions (the black area in the lower panel in Figure 5C) is substantially shifted in the negative direction, *r*_z_ *mean* = .463, *sem* = .125, *t*_17_ = 3.697, *p*=.0018; note that the shape of this distribution, including the few outliers in the positive direction, is similar to the corresponding distribution obtained in Experiment 1 (the lower panel in Figure 3C). The distribution for the relationship for the spatial-average and control conditions (the green area in the lower panel in Figure 5C) is also shifted in the negative direction, *r*_z_ *mean* = .200, *sem* = 0.078, *t*_17_ = 2.561, *p*=.02, but to a significantly lesser extent, *r*_z_ *mean difference* = .262, *sem* = 0.123, *t*_17_ = 2.131, *p*=.048.

Overall, we have replicated Experiment 1. The association between posterior beta-power reductions and higher aesthetic ratings as a function of increasing degree of multistability is virtually identical in the two experiments (Figures 3 and 5). A similar but weaker association was obtained for the spatial-averaged condition (Figure 5C), consistent with the interpretation that the spatially averaged versions that approximate the output from the low-spatial-frequency channels might have indirectly generated multistability-induced effects through learned associations. Finally, the fact that the multistable condition substantially reduced posterior beta power relative to the local-control, global-control, and spatial-average conditions (Figures 3A and 5A) strongly suggests that posterior beta power reductions are driven by spatiotemporal multistability over and above any inherent temporal dynamics.

## General discussion

Spatially heterogeneous—spatiotemporally multistable—flicker reduced posterior EEG beta power (~11Hz to ~23Hz in Experiment 1 and ~14Hz to ~20Hz in Experiment 2) relative to the spatially homogeneous—monostable—flicker controls that were matched in the temporal dynamics of local luminance changes, luminance changes at any location, and spatially averaged luminance changes. Even the lowest degree of multistability that included only two independently flickering regions substantially reduced posterior beta power relative to the monostable controls, while the highest degree of multistability with sixteen independently flickering regions maximally reduced posterior beta power. The fact that these effects were localized to the posterior sites and were equivalent with or without the accompanying (annoying) click sounds suggest that these beta-power reductions reflect the visual processing of spatially heterogeneous flicker rather than higher-order effects such as relaxation or reduced arousal from viewing flicker that may resemble nature.

Strictly based on the data, our EEG results demonstrate that viewing displays containing a larger number of independently flickering regions induces a larger reduction in posterior beta power. A larger number of independently flickering regions provides a greater spatiotemporal variety in motion energy, which in turn affords a greater variety of potential motion interpretations, resulting in a greater degree of spatiotemporal multistability in motion perception. Therefore, our preferred interpretation is that the processing of spatiotemporal multistability reduces posterior beta power.

Our behavioral results suggest that people also aesthetically prefer flicker with greater degrees of spatiotemporal multistability. Note that the negative effects of the click sounds on aesthetic ratings in both experiments were subtractive and did not interact with the flicker conditions, suggesting that the auditory influences additively contributed at the decision stage. If the observers responded based on their general impressions such as their moods or feelings evoked by the supramodal stimulus experience (e.g., being reminded of nature), the auditory influences would have likely been nonlinear. Thus, our preferred interpretation is that the ratings reflect the observers’ aesthetic responses to the visual experience of multistable flicker.

A potential objection to these interpretations might be that beta-power reductions and aesthetic preferences could both have been induced by complexity rather than multistability. Certainly, our stimuli with 16 independently flickering regions appeared “more complex” than those with 2 independently flickering regions. However, this is largely a semantic issue. A visual stimulus affording a greater degree of multistability necessarily has to appear more complex in the sense that a straightforward definition of perceptual complexity of a visual stimulus is the number of perceptual interpretations it affords; that is, a stimulus that can be interpreted in many different ways is more complex than a stimulus that can be interpreted in few ways or only one way.

A more substantive concern with our interpretation that beta-power reductions and aesthetic preferences were driven by the processing of spatiotemporal multistability is that these effects might have been elicited by some other variables that covaried with our manipulation of spatiotemporal multistability, that is, with the number of independently flickering regions. Temporal statistics covaried with the number of independently flickering regions. We controlled for their potential influences by including the local-control, global-control, and spatial-averaged conditions. In particular, neither the beta-power reductions nor aesthetic preferences associated with flicker multistability could be due to increased temporal density of events (matched by the global-control condition) or increased temporal smoothness due to internal spatial summation (matched by the spatial-average condition). Spatial frequency also covaried with the number of flickering regions; a flicker stimulus with a greater number of independently flickering regions contained higher spatial-frequency components. To alleviate potential spatial-frequency effects, we presented all flicker stimuli on the pedestals of 16 rectangles (Figure 1). Nevertheless, potential roles of higher spatial-frequency components in beta-power reduction and/or increased aesthetic preference would be an empirical question. Frund et al. (2007) examined evoked spectral powers in the alpha (8-13Hz) and gamma (30-85Hz) bands as a function of spatial frequency using Gabors (unfortunately, no results are reported for the beta band), showing that alpha power increased (while gamma power decreased) with increasing spatial frequency. While we obtained robust multistability-based power reductions in the beta range (Figures 2B and 4B), the multistable conditions also numerically reduced power in the neighboring alpha band in spite of the higher spatial-frequency components. This is the opposite of Frund et al.’s results, suggesting that our beta-power reduction result was not driven by spatial frequency (though this comparison needs to be taken with a grain of salt because Frund et al. primarily used single-frequency Gabors and their results with a compound Gabor suggested non-linear effects of spatial frequency components). Further, aesthetic preferences for static images depend on overall spatial-frequency composition (e.g., Vannucci, Gori, & Kojima, 2014); if anything, increased power in higher-frequency components (within the range relevant to our stimuli) causes visual discomfort (Fernandez & Wilkins, 2008), suggesting that the increased preferences we obtained for the spatiotemporally multistable stimuli could not be due to their increased higher spatial-frequency components. These considerations reasonably rule out alternative interpretations of our results based on temporal statistics or spatial clutter that covaried with our manipulation of spatiotemporal multistabilty.

The core result from the current study is that posterior beta-power reductions and aesthetic preferences are both closely associated with the increased degree of spatiotemporal multistability of flicker. The result is robust as the two experiments yielded virtually identical relationships. As briefly described in the introduction section, our results may provide a converging perspective on the literature on beta-power reductions and visual motion perception to support a broad hypothesis; that is, natural phenomena such as flickering flames, rippling water surfaces, rustling leaves, and spattering rain drops are pleasant to the senses potentially because they present spatiotemporally heterogeneous flicker that affords spatiotemporal multistability that calibrates visual motion processing. In the remaining paragraphs, we elaborate on this perspective.

Posterior/central MEG alpha/beta-power reductions have been observed during visual tasks with lower cognitive and/or attentional demands (e.g., Donner & Siegel, 2011; Donner et al., 2007; Siegel et al., 2008). MEG/EEG beta-power reductions have also been reported during the perception of local feature motions as opposed to global object motions (Aissani et al., 2014). Because top-down controls and perceptual grouping require long-range neural interactions, alpha/beta-power reductions accompanying processes that require less attentive scrutiny or grouping is consistent with the idea that the amount of posterior/central alpha/beta power reflects the amount of engagement of top-down long-range neural interactions in visual processing (e.g., Donner & Siegel, 2011). This interpretation is also consistent with the evidence suggesting that feedback neural interactions are mediated by alpha/beta bands (10Hz–20Hz; Bastos et al., 2015; Michalareas et al., 2016). Nevertheless, posterior/central alpha/beta reductions associated with visual competition and the perception of biologically plausible motions do not readily fit into this framework. Posterior EEG beta power was reduced during binocular rivalry and illusory motion reversals (Piantoni et al., 2010). The authors postulated that the reduced beta power reflected the reduced size of the neural population synchronously representing the visible percept; the population would be largest when a single percept dominates (as in typical visual experiences), reduced during binocular rivalry due to the mutual inhibition of the populations representing the competing percepts, and much reduced during the infrequent perception of illusory reversed motions due to the strong inhibition from the large population representing the veridical motion. Posterior/central EEG alpha/beta power was also reduced during the perception of biologically plausible motions (Meirovitch et al., 2015). The authors postulated that the alpha/beta reductions reflected an engagement of the prioritized motion mechanisms that mediated action-perception coupling. While alpha/beta reductions may reflect multiple different processes, a common theoretical framework that may potentially accommodate all these examples is that posterior/central alpha/beta-power reductions, indicative of reduced top-down long-range neural interactions, are associated with the processing of sensory signals that conform to the biases (expectations) of the visual system.

More top-down scrutiny would be required when the bottom-up visual processing does not readily provide task relevant information; conversely, less top-down scrutiny would be required when visual processing is allowed to proceed according to the current biases (states) of the visual system. During binocular rivalry, the perceived image spontaneously alternates between the competing images presented to the two eyes, while perceptual alternations occur when the processing of the suppressed image overcomes the processing of the dominant image based on neural adaptation and noise (e.g., Kim et al., 2006; Wilson, 2007; Alais et al., 2010), that is, when the state of the visual system becomes more biased in favor of processing the suppressed image. In the illusory-motion-reversal displays, the dominant percept of veridical motion occasionally reverses when the motion detectors tuned to the veridical motion sufficiently weaken due to adaptation, that is, when the state of the visual system becomes strongly biased in favor of processing the reversed motion. Evidence suggests that the visual system prioritizes the processing of biologically plausible motions (e.g., Allison et al., 2000; Grossman et al., 2000; Plass et al., 2014), that is, the visual motion mechanisms may be biased in favor of processing motion signals that conform to familiar biological constraints. This interpretation is consistent with the finding that motion-perception related alpha/beta-power reductions were greater for individuals with greater familiarity with the observed movements (Orgs et al., 2008). Thus, most examples of alpha/beta-power reductions associated with visual processing could potentially be explained by postulating that the processing of sensory signals that conform to the current biases (expectations) of the visual system imposes reduced demands for long-range neural interactions, which is reflected in EEG and MEG recordings as reduced posterior/central alpha/beta power.

How might spatially heterogeneous flicker provide visual signals that conform to the current biases of the visual system? Spatially heterogeneous flicker generates spatiotemporally multistable visual signals that simultaneously contain motion energies (e.g., Adelson and Bergen, 1985; Georgeson and Scott-Samuel, 1999) in multiple directions, speeds, and spatial scales. Spatially heterogeneous flicker thus provides similarly effective input to competing motion detectors tuned to a variety of directions, speeds, and optic-flow patterns such as expansion/contraction and rotation in multiple processing stages (see Orban, 2008, for a review). As a consequence, spatially heterogeneous flicker may allow visual motion processes to transpire according to the current sensitivity asymmetries among motion detectors. For example, if the upward-direction (or clockwise-rotation, etc.) tuned motion detectors happened to be more sensitive than those tuned to other spatiotemporal patterns in a given region at a given time, the omnidirectional motion energies in multistable-flicker signals would allow the most sensitive upward-direction (or clockwise-rotation, etc.) tuned detectors to win the competition and be activated. In this sense, the current result of posterior beta-power reductions in response to spatially heterogeneous flicker is consistent with the interpretation that multistable flicker engages visual processes that conform to the current biases of the visual system, thereby engaging reduced top-down processes involving long-range neural interactions.

The fact that multistable flicker activates motion detectors according to their sensitivity differences suggests that it may have a restorative effect. Selective activation of motion detectors with greater sensitivity would calibrate the system-wide spatiotemporal sensitivity by selectively adapting the detectors with elevated sensitivities. It is plausible that such sensory calibration may be perceptually beneficial so that multistable flicker that facilitates it may engage the reward system. Consistent with this line of reasoning, multistable flicker was aesthetically preferred in the current study. In both experiments, flicker with the highest degree of multistability (with sixteen independent flickering regions) was strongly preferred, yielding the average aesthetic rating of ~3.5 (leftmost panels in Figures 3B and 5B) with 4 indicating maximum preference and 2.5 indicating neutral. The 2-region and 4-region flicker primarily contained horizontal, vertical, and rotational motion signals at a relatively large scale (with a quadrant as a spatial unit), whereas the 16-region flicker additionally contained motion signals in intermediate directions as well as a greater variety of motion patterns (e.g., rotation, expansion, contraction, etc.) in multiple scales. Thus, the 16-region flicker should have been more effective at generating activation patterns closely conforming to the spatiotemporal sensitivity biases of motion detectors and therefore more effective at neutralizing motion detector sensitivity imbalances than the 2- and 4-region flicker, while the 1-region flicker should have been ineffective. Consistent with this interpretation, while the 2- and 4-region flicker produced substantial but similar degrees of posterior beta-power reductions and aesthetic preferences relative to the 1-region flicker, the 16-region flicker produced much larger effects. We confirmed that these associations were consistent across observers in both experiments. Thus, our overall results suggest that the ability of spatially heterogeneous flicker to adaptively calibrate motion detectors by engaging visual processes conforming to the current sensory biases—indexed by the multistability-dependent posterior beta-power reductions—is closely associated with aesthetic responses to flicker.

In conclusion, the current results are consistent with an overarching interpretation that naturally abundant spatially heterogeneous flicker elicits aesthetic responses from purely dynamic information based on its spatiotemporal multistability that contributes to sensory sensitivity calibration. We acknowledge that this overarching interpretation is speculative. For example, future research may discover that the obtained relationship between posterior beta-power reductions and aesthetic responses may be driven by spatiotemporal factors that covaried with our manipulation of spatiotemporal multistability (we considered temporal statistics and spatial frequency, but there may be others). Further, although substantial evidence (primarily in the literature on flicker motion aftereffects) suggests that observing spatiotemporally multistable stimuli reduces motion sensitivity biases, future research needs to investigate how (or whether) this type of sensory calibration improves motion perception. Notwithstanding the need for these and other future investigations, we believe that it is worth disseminating the current results that may inspire a general hypothesis that spatiotemporal multistability may be a core principle underlying the neural, functional, and aesthetic impacts of experiencing visual dynamics in nature. Finally, our results extend the postulated link between posterior/central alpha/beta-band power reductions and reduced top-down controls in the context of experiencing multistable sensory stimulation.

